# Examining excitation-inhibition modulation during social processing using functional Magnetic Resonance Spectroscopy

**DOI:** 10.64898/2025.12.03.691778

**Authors:** Talitha C. Ford, Duanghathai Pasanta, Nina-Francesca Parrella, Peter G. Enticott, David J. White, Nicolaas A. Puts

## Abstract

An imbalance of brain excitation-inhibition has been implicated in social communication function across neurodevelopmental and psychiatric conditions. Recent advances in functional magnetic resonance spectroscopy (fMRS) allow non-invasive measurement of dynamic changes in excitation-inhibition neurometabolites (glutamate and GABA), revealing potential links with cognitive processes. This study used functional MRS (fMRS) to investigate whether socially-laden stimuli modulate excitatory or inhibitory neurometabolites, and whether modulation relates to social function.

Forty neurotypical adults (18-40 years; 20 male) underwent 3T MRI, with fMRS data acquired using a J-difference edited MRS sequence. GABA+macromolecules (GABA+) and glutamate+glutamine (Glx) were quantified in a right social brain region (superior temporal/temporoparietal junction [STG/TPJ]) and a control region (occipital cortex).

Participants viewed a fixation cross (traditional resting state MRS; 5 min), silent video clips of social interaction, and a flashing checkerboard as control stimulus (5 min each; counterbalanced). The Social Responsiveness Scale (SRS-2) assessed social function.

Block analysis showed no significant differences between resting and functional (social or checkerboard) GABA+ or Glx levels (*p*s<.05), although Bayesian analyses showed evidence for decreased STG/TPJ GABA+ and increased occipital cortex Glx for checkerboard compared to rest. Higher SRS-2 scores were associated with higher resting STG/TPJ Glx but not GABA+, and functional metabolite levels were not associated with SRS-2 scores (*p*s>.1). Sliding time- window analyses showed significantly reduced Glx in social brain during social stimulation.

Despite sound data collection and analysis techniques, shifts in Glx and GABA+ following social stimuli were minimal. Future research could utilise more challenging social stimuli and examine alternate social brain regions.

## 1. Introduction

Social information processing difficulties are central to several neurodevelopmental and psychiatric conditions, such as autism spectrum disorder, social anxiety disorder, attention deficit-hyperactivity disorder, and schizophrenia spectrum disorders (Aydin et al., 2023; Chisholm et al., 2015; Clark et al., 1999; Oliver et al., 2021). Several lines of evidence suggest that an imbalance in neural excitation and/or inhibition underlies these social information processing difficulties (Alabdali et al., 2014; Canitano & Palumbi, 2021; Ford et al., 2017b; Ford & Crewther, 2016; Mamiya et al., 2021; Parrella et al., 2024; Rubenstein & Merzenich, 2003; Yizhar et al., 2011). A balance in neural excitation and suppression is critical for stable perception, cognition, and behaviour (Donahue et al., 2010; Puts et al., 2011; Stagg, 2014; Yoon et al., 2010). Thus, an imbalance in excitatory or inhibitory processes has direct implications (Kolasinski et al., 2019; Stagg, 2014) for interpreting social information in the environment. Non-invasive neuroscientific techniques, such as magnetic resonance spectroscopy (MRS), can quantify neurometabolites that are directly involved in brain excitation (glutamate) and inhibition (GABA). MRS studies have shown that reduced social processing functionality is associated with increased glutamate levels in anterior cingulate cortex (ACC), superior temporal regions, and occipital cortex (Cochran et al., 2015; Ford et al., 2017a; Sousa et al., 2025) and reduced GABA levels in ACC, superior temporal regions, medial prefrontal cortex, and occipital cortex (Brix et al., 2015; Carvalho Pereira et al., 2018; Cochran et al., 2015; Ford et al., 2017a; Kirkovski et al., 2018; Sousa et al., 2025).

Traditional human MRS methods only allow for the quantification of regional brain metabolite levels at rest (e.g., while viewing a fixation cross), which limits understanding of how metabolites are involved in real-time cognitive processes. Recent advances in MRS data collection and analysis techniques have led to the ability to examine *dynamic* metabolite levels; that is, during engagement with sensory or cognitive stimulation (Pasanta et al., 2023). During stimulation, there is an increase in cellular glutamate and GABA release and recycling, which may result in a relative change in the proportions of these neurotransmitters (Ip et al., 2017; Lin et al., 2012). Functional MRS (fMRS) studies have shown increases in excitatory glutamate when participants: view a flashing checkerboard (Bednařík et al., 2015; Ip et al., 2017, 2019; Koush et al., 2021; Lynn et al., 2018) or visual contrast stimulus (Lin et al., 2012); view novel stimuli in a repetition suppression task (Apšvalka et al., 2015); engage in motor tasks (Lynn et al., 2018; Schaller et al., 2014); engage in cognitive tasks (Craven et al., 2023; Kühn et al., 2016; Taylor et al., 2015; Woodcock et al., 2018) and task learning (Stanley et al., 2017); and engage in mental imagery (Huang et al., 2015). Although there are fewer studies examining functional GABA, shifts in GABA levels have been reported during visual stimulation (Koush et al., 2021; Lin et al., 2012), and increased GABA has been reported during cognitive task (Kühn et al., 2016). However, there is substantial variability between study methods and subsequent findings (Pasanta et al., 2023), with no change in insular glutamate or GABA reported in response to pain stimuli (Cleve et al., 2017), no change in medial ACC GABA during cognitive tasks, and no change in primary visual cortex GABA during flashing checkerboard (Ip et al., 2019). Nevertheless, fMRS has potential to better understand brain-behaviour relationships.

Functional MRI (fMRI) is well established as a tool for examining neural activity in brain regions associated with sensation, perception, and cognition through measurement of the blood oxygenation level-dependent (BOLD) signal (Logothetis et al., 2001). BOLD signal is interpreted as an increase in regional energy consumption, through the interplay between cerebral blood flow, cerebral blood volume, and cerebral metabolic rate of oxygen consumption (Donahue et al., 2010; Logothetis et al., 2001). The relationship between BOLD signal and regional glutamate and GABA levels is less understood, although glutamate and GABA play a significant role during activity-related energy metabolism in the brain (Ip et al., 2017, 2019). Reduced baseline GABA levels in visual cortex have been associated with increased BOLD signal in visual cortex during visual stimulation (Donahue et al., 2010; Muthukumaraswamy et al., 2009), while glutamate levels in visual cortex have been shown to increase with increased BOLD signal during visual stimulation (Bednařík et al., 2015).

Increased glutamate in dorsolateral prefrontal cortex (Vijayakumari et al., 2018) and reduced GABA in the anterior cingulate (Kühn et al., 2016) have also been associated with increased BOLD signal during cognitive tasks. No studies to date have examined the relationship between BOLD signal change and shift in GABA and glutamate during social information processing. As such, a social information processing task that has been shown to elicit BOLD signal in right superior temporal regions was utilised herein (Deen et al., 2015), with the MRS voxel location centred around the expected BOLD signal location.

To date, no study has comprehensively investigated the extent to which fMRS can detect changes in excitatory glutamate and inhibitory GABA during social information processing, or how such shifts in relate to social communication function. This study therefore examined changes in Glx (glutamate + glutamine) and GABA+ (GABA co-edited with macromolecules) levels while participants were engaged in a social task. The social task and MRS voxel location were selected based on evidence that the right superior temporal gyrus (STG) and temporoparietal junction (TPJ) shows robust and selective responses to dynamic social stimuli, relative to non-social visual input (Deen et al., 2015; Pitcher et al., 2011). This functional specificity makes STG/TPJ an optimal target for detecting task-related modulation of excitation-inhibition balance, while a control voxel in occipital cortex was included to index general visual processing. It was hypothesised that viewing dynamic social stimuli would modulate Glx and GABA+ levels in a right social brain voxel including STG/TPJ and insula cortex, but not in a control voxel in occipital cortex. Similarly, it was hypothesised that visual stimulation via flashing checkerboard would lead to a shift in Glx and GABA+ in occipital cortex, but not social brain. We also probed the relationship between resting state and dynamic social brain Glx and/or GABA+ and social communication function via the Social Responsiveness Scale (SRS-2), a gold standard Research Domain Criteria (RDoC) *Social Processes* measure (Constantino & Gruber, 2005; Cuthbert & Insel, 2013; Insel, 2010).

Finally, we examined the relationship between fMRI BOLD signal and dynamic Glx and GABA+ levels in social brain and occipital cortex during dynamic social and visual checkerboard stimuli, respectively.

## 2. Methods

The study was approved by Swinburne University of Technology’s Human Research Ethics Committee (20202614-3319) in accordance with the National Statement of Human Research and the Declaration of Helsinki. The study was pre-registered on OSF (https://osf.io/y7jqp). Written informed consent was obtained from all participants prior to undergoing study procedures.

### 2.1. Participants

Forty neurotypical young adults were recruited for this study (20 male, age 18-40 years). All participants self-reported fluency in written and spoken English, normal or corrected-to-normal vision, normal hearing, and no intellectual disability. Participants did not have any psychiatric, developmental, substance use, genetic, and neurological conditions (based on self-report). At the time of testing, participants were free from psychoactive medications (>1 month), nicotine (abstained >12 hrs), caffeine (abstained >12 hrs), alcohol (abstained >12 hrs), and recreational drugs (abstained >1 week). All participants met Swinburne Neuroimaging MRI safety criteria, including that female participants were not pregnant or lactating (self-reported). Due to the use of a flashing checkerboard stimulus in the task protocol, participants were also screened for sensitivity to the stimulus (e.g., photosensitivity, causing migraine); no participants were excluded due to sensitivity).

Sample size was determined *a priori* using a repeated-measures ANOVA framework (*f* = 0.2, *α* = 0.05, power = 0.8), with the effect size selected based on conventional small-to-moderate benchmarks (Cohen, 1988) given the limited prior fMRS data and modest effect sizes typically reported, to detect changes in GABA+ and Glx levels across resting state and functional blocks (*n* = 36). A total of 40 participants were recruited to allow for potential exclusions due to poor spectra quality. Although a repeated measures ANOVA was originally planned, the primary hypothesis was tested using a linear mixed-effects models to account for missing data and participant-level random effects. The planned sample size remained appropriate given that mixed-effects models have comparable or greater statistical efficiency relative to repeated measures ANOVA (Muhammad, 2023).

Participants completed the Social Responsiveness Scale, 2^nd^ Edition (SRS-2) to assess social communication function (Constantino & Gruber, 2005) (RDoC Social Processes domain). The SRS-2 is a 65-item measure of symptom severity across five autism-related categories: Social Awareness (8 items), Social Motivation (11 items), Social Cognition (12 items), Social Communication (22 items), and Repetitive Behaviors/Restricted Interests (12 items).

Responses are measured on a 4-point scale from 0 (not true) to 3 (almost always true). The SRS-2 raw scores were utilised herein rather than converted *t*-scores in order to preserve variability in our non-clinical population (score range 7-102).

### 2.2. Magnetic Resonance Protocol

MRI and MRS data were collected using a 3T Siemens whole-body Prisma scanner (Siemens, Erlangen, Germany) with 32-channel head coil at Swinburne Neuroimaging. Participants underwent two 60-minute MRI scans, which were separated by a break of 30-90 minutes. For each scan, T1-weighted structural images were acquired sagittally using a magnetisation pre-prepared rapid gradient echo (MPRAGE) pulse sequence with an inversion recovery (repetition time [TR] = 1900ms, echo time [TE] = 2.34ms, inversion time [TI] = 900ms, flip angle α = 9°, field of view [FOV] = 256mm, 192 sagittal slices, acceleration factor = 3, isotropic voxel size = .89mm^3^, bandwidth = 200Hz/Px, acquisition time = 3:51mins).

Structural images were used for MRS voxel placement, tissue composition correction, and co-registration to fMRI BOLD data.

#### 2.2.1. MRS Protocol

All resting state and functional MRS data were collected in one continuous run using a Hadamard Encoding and Reconstruction of Mescher-Garwood-Edited Spectroscopy (HERMES) sequence (Mescher et al., 1998; Saleh et al., 2020), an editing sequence that allows for isolated GABA+ quantification (TR = 2000 ms, TE = 80 ms, 780 transients, acquisition time = 26:08 mins). Editing pulses were placed at 1.90 (edit-ON1), 4.56 (edit- ON2), and 7.50 (edit-OFF) ppm. Given the HERMES editing sequence co-edits macromolecular signals at 3ppm we refer to GABA+. During the first scan, GABA+ and Glx levels were quantified in a voxel positioned in the right “social brain” region including TPJ, STG, and insular cortex (40 x 25 x 25 mm voxel, see Figure 1), which are strongly implicated in social information processing (Blakemore, 2008; Jung et al., 2019; Scrivener & Reader, 2022). During the second scan, GABA+ and Glx levels were quantified in a control region predominantly within visual cortex (30 x 30 x 30 mm voxel, see Figure 1) selected to index lower-level visual processing relative to the socially selective responses observed in STG/TPJ (Deen et al., 2015; Pasanta et al., 2024; Pitcher et al., 2011). Due to anatomical and shimming constraints associated with placement of a large, edited MRS voxel in visual cortex, the final voxel placement extended beyond the intended primary visual cortex (V1) region into secondary visual cortex (V2). A matched acquisition without water suppression was acquired before and after each HERMES sequence for eddy-current correction and quantification (8 averages) (Edden et al., 2016).

**Figure 1.**
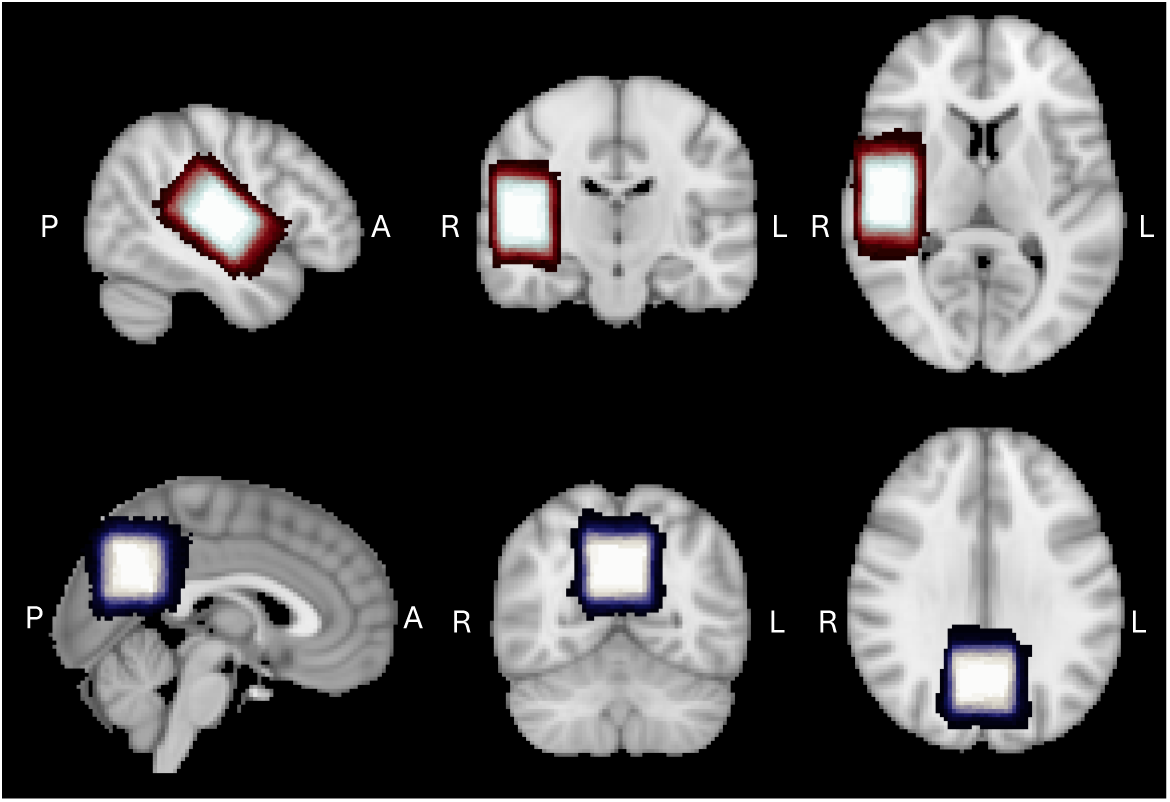
Average magnetic resonance spectroscopy (MRS) voxel placement. Top: Group average placement of the right social brain voxel including superior temporal gyrus and temporoparietal junction. Bottom: Group average placement of the control voxel in visual cortex. MRS voxel masks warped to MNI space.

The fMRS task was a modified version of that published previously (Deen et al., 2015; Pitcher et al., 2011) and our own feasibility study (Pasanta et al., 2024). Stimuli consisted of a white fixation cross on a black background (resting state), short 3-second movies of children engaging in social interaction (dynamic faces; 100 videos per stimulus block) (Deen et al., 2015; Pitcher et al., 2011), and a 7Hz blue and yellow flashing checkerboard. Figure 2 illustrates the stimuli block order and relevant HERMES sequence parameters. Block order and timings were as follows: 1) resting state block (fixation cross1; 5 mins, 150 TRs), 2) functional block (dynamic faces or checkerboard, counterbalanced; 5 mins, 150 TRs), 3) resting state block (fixation cross2 8 mins, 240 TRs), 4) functional block (opposite to that presented in first stimulus block; 5 mins, 150 TRs), 5) resting state block (fixation cross3; 3 mins, 90 TRs). Between each block was a 5-second (2.5 TR) text display noting the end of the current block and reminder to remain still for the following block. Participants were instructed to keep their eyes open and passively view the screen for the duration of the sequence.

**Figure 2.**
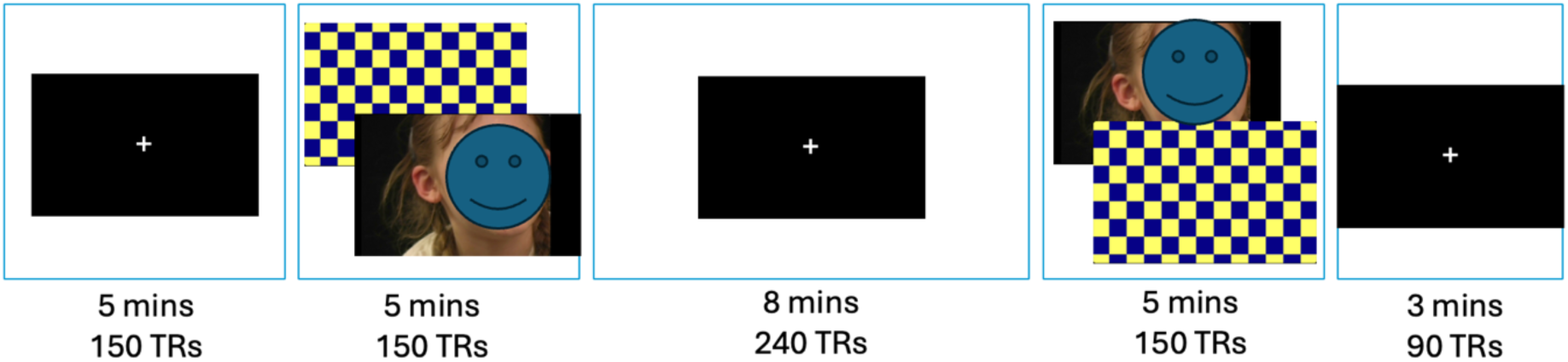
fMRS stimulus presentation timing. From left to right, rest (fixation cross, 5 mins, 150 transients), functional block of 7Hz flashing checkerboard or dynamic faces (100 x 3-sec movies; 5 mins, 150 transients), rest (8 mins, 240 transients), functional block (5 mins, 150 transients), rest (3 mins, 90 transients). Note: face image de-identified.

#### 2.2.2. fMRI Protocol

fMRI data were collected immediately following the HERMES sequence in the second MRI scan only to investigate the relationship between stimulus-related metabolite levels and BOLD signal. The fMRI scanner protocol was as follows: T2*-weighted echo planar imaging (EPI) pulse sequence sensitive to blood oxygen level–dependent (BOLD) contrast (TR = 2500ms, TE = 30ms, flip angle α = 90°, FOV = 192mm, matrix = 64×64, slice thickness = 3mm, voxel size 3mm^3^, slice gap = 0.6mm, 46 axial slices, acquisition time = 5:34 mins). In line with the fMRS task, the fMRI task was a modified version of that published previously (Deen et al., 2015; Pitcher et al., 2011). Participants passively viewed two runs of six stimulus blocks consisting of two blocks each of dynamic faces, dynamic objects, and the flashing checkerboard (counterbalanced). An 18 sec fixation cross was presented at the beginning and end of each block combination, with a 10 second rest between each run. Blocks were presented in a palindromic fashion in each run (e.g., + F C O + O C F + rest + C F O + O F C +, see Supplementary Figure 1). The dynamic faces stimuli have been shown to elicit strong BOLD signal in right superior temporal regions (Deen et al., 2015; Pitcher et al., 2011), while the flashing checkerboard was included as a control visual stimulus. The objects condition is beyond the scope of this paper and is thus not presented here. Stimulus blocks ran for 18 seconds each (6 x 3-sec movies of faces or objects, 18-sec checkerboard).

### 2.3. Data Analysis

#### 2.3.1. MRS metabolite quantification

All MRS data were analysed using Gannet version 3.3.2 with MATLAB, a purpose-built software package for quantifying edited GABA spectra (Edden et al., 2014) with custom-code for functional analysis. First, the effect of scanner drift over the entire acquisition (780 transients) was visually inspected for each participant based on Cr frequency trace generated by GannetLoad after SpecReg HERMES was applied – there was no evidence of meaningful scanner drift for any participant. Separate block analyses were then conducted for the first resting state block (fixation cross1; TRs 1-152) and the two functional blocks (TRs 153-304 and 547-698). For each block, data were pre-processed and fit using multi-step frequency and phase correction designed for HERMES data (Mikkelsen et al., 2018).

Individual MRS voxels were then co-registered to the participant’s T1 structural image using SPM12 (www.fil.ion.ucl.ac.uk/spm), and tissue segmentation was conducted to calculate ratios of grey matter (GM), white matter (WM), and cerebrospinal fluid (CSF) (Ashburner & Friston, 2005). Finally, metabolites were quantified and corrected for GM, WM, and CSF using the Harris method (Harris et al., 2015).

As an exploratory approach, we analysed spectra using a sliding time-window to achieve higher temporal resolution of metabolite changes (Pasanta et al., 2024). A fixed window of 60 TRs was applied, advancing in steps of 10 TRs, and repeated until the end of the acquisition. This resulted in 73 timepoints per participant per voxel; total N = 2,920 time- points per voxel per acquisition. Data-points where conditions overlapped were labelled accordingly. For example, cross+checkerboard indicates that the analysis window contained both the fixation cross (resting state) and checkerboard presentation, while cross+faces denote overlap between the fixation cross and face stimuli. This approach yielded an effective temporal resolution of 1 minute and 15 seconds, with each data-point representing 75 seconds of data (Pasanta et al., 2024).

#### 2.3.2. fMRI BOLD data analysis

Preprocessing was performed using fMRIPrep 25.1.3 (Esteban et al., 2018, 2019), which is based on Nipype 1.10.0 (Gorgolewski et al., 2011, 2018). Please see Supplementary Material for the complete fMRIPrep pipeline. Volume-based spatial normalization to the MNI152NLin6Asym standard spaces was performed through nonlinear registration with antsRegistration (ANTs 2.6.2), using brain-extracted versions of both T1w reference and the T1w template. This normalised dataset was entered into FSL FEAT for level 1 data processing (FMRI Expert Analysis Tool) Version 6.00, part of FSL (FMRIB’s Software Library, www.fmrib.ox.ac.uk/fsl)(Jenkinson et al., 2012).

The following pre-statistics processing was applied in FEAT: spatial smoothing using a Gaussian kernel of FWHM 5mm; grand-mean intensity normalisation of the entire 4D dataset by a single multiplicative factor; highpass temporal filtering at 72.0 sec (Gaussian-weighted least-squares straight line fitting, with sigma=36.0s). Time-series statistical analysis was carried out using FILM with local autocorrelation correction (Woolrich et al., 2001).

MRS voxel masks (social brain and control region) were created for each participant using ANTS antsApplyTransforms and mean BOLD signal (stimulus minus rest) from each participant voxel mask was extracted using FSL featquery to investigate associations with GABA+ and Glx levels.

### 2.4. Statistical analysis

Statistical analyses were conducted in R (version 2025.05.0). There were no age or sex effects on GABA+ or Glx levels (*p*s > .05). There was, however, an effect of block order for the social brain voxel data (*p*s < .05), therefore block order was entered as a covariate in subsequent analyses. To test the primary hypothesis, that dynamic social stimuli would result in a shift in Glx and GABA+ concentration in a right social brain voxel, two separate mixed-effects models were designed using the *lme4* package in R (Bates et al., 2015), with GABA+ or Glx entered as the outcome variable. All assumptions for mixed effects models were met, except for normality of residuals for GABA+ (skew = 1.16), which was subsequently log transformed. For both models, voxel (social brain, control), stimulus (fixation cross, faces, checkerboard), and block order were entered as fixed effects, and participant was entered as the random effect (see Supplementary Material for full model design). Degrees of freedom for fixed effects were estimated using the Satterthwaite approximation, as implemented in the lmerTest package, reflecting the number of observations and model structure rather than the number of participants alone.

Follow-up Bayesian analyses were then conducted using the *brms* and *BayesFactor* packages in R (Bürkner, 2017, 2018, 2021; Morey & Rouder, 2024). The ratio of Glx and GABA+ was calculated by dividing Glx by GABA+ and exploratory analyses we conducted as per the above. Results for Glx/GABA ratios are presented in the Supplementary Material.

For sliding time-window analysis, an additional autoregressive (AR) term was used to account for autocorrelation between data-points to assess the effect of stimulus type on metabolite changes where the first resting state block was used as a baseline. Weakly informative priors were chosen for Bayesian analyses of sliding time-window. The model choice for sliding time-window Bayesian model was validated through model comparison using Pareto-Smoothed Importance Sampling Leave-One-Out Cross-Validation (PSIS-LOO). Results showed that the additional autoregressive terms substantially improved the model in comparison to the model without autoregressive term for all metabolites. Models were estimated in *brms* using Hamiltonian Monte Carlo. Four chains of iterations each (1,000 warm-ups) yielded 8,000 post–warm-up draws. All parameters had R-hat <1.01, indicating good convergence.

The Range of Practical Equivalence (ROPE) test was used to determine whether the effect could be considered practically significant (Cohen, 1992; Kruschke, 2018). Specifically, we examined whether the 89% Highest Density Interval (HDI) of the posterior distribution fell entirely within the ROPE; defined as -0.1 to 0.1 times the standard deviation of the outcome variable (Kruschke, 2015, 2018). Analyses were conducted in R using the *bayestestR* package (Makowski et al., 2025).

To examine the relationship between resting state and functional GABA+ and Glx levels and SRS-2 scores, Spearman’s rho coefficients were calculated due to moderate-substantial skew in social brain GABA+ levels. FDR was controlled for within each metabolite combination using the Benjamini-Hochberg method (Benjamini & Hochberg, 1995). Exploratory correlation analyses were conducted for the relationship between Glx/GABA+ ratios and SRS scores, with results presented in the Supplementary Material.

Finally, the relationship between relative BOLD signal (stimulus minus baseline) and relative GABA+ and Glx levels (stimulus minus resting state) was examined for the faces and checkerboard stimuli. BOLD signal was extracted from the MRS mask region of interest at the individual participant level. Pearsons correlations were conducted controlling for FDR within each metabolite using the Benjamini-Hochberg method (Benjamini & Hochberg, 1995).

## 3. Results

Participant characteristics are presented in Table 1. There were no significant sex differences in age, handedness, years of formal education, or history of psychiatric conditions such as depression or anxiety. There were also no sex differences in SRS-2.

**Table 1.** Total participant sample characteristics.

|  | Total | Female | Male | Sex differences |
| --- | --- | --- | --- | --- |
| N | 40 | 20 | 20 |  |
| Age (M(SD)) | 30.1(5.8) | 30.4(6.3) | 29.7(5.5) | $t(37)=.38, p=.710$ |
| Handedness (L:R:A) | 5:33:2 | 2:18:0 | 3:15:2 | $\chi^2(2)=2.5, p=.290$ |
| Years formal education (M(SD)) | 17.4(4.1) | 16.7(3.9) | 18.1(4.2) | $t(37)=-1.0, p=.322$ |
| Psychiatric history (N(%)) | 6(15%) | 3(15%) | 3(15%) | $\chi^2(2)=1.0, p=.597$ |
| SRS-2 total (M(SD)) | 42.8(24.0) | 36.2(19.4) | 49.4(26.7) | $t(34.6)=-1.8, p=.081$ |

Notes: M = mean, SD = standard deviation, L = left; R = right; A = ambidextrous; SRS-2 = Social Responsiveness Scale – 2<sup>nd</sup> Edition.

### 3.1. MRS data quality

MRS spectra quality was assessed for each block (*n* = 3) within each voxel (*n* = 2) for each participant (*n* = 40, total 240 datasets), with individual datasets removed from further analysis where water or NAA FWHM > 15Hz, GABA+ or Glx fit error > 15%, GABA+ signal-to- noise ratio (SNR) < 5, or Glx SNR < 10. These thresholds were defined *a priori* based on commonly used criteria in edited MRS studies and established methodological recommendations (Mullins et al., 2014; Near et al., 2021). This resulted in 8 datasets excluded (water FWHM = 3, GABA+ fit error = 4, Glx SNR = 1; 96.67% retained). An additional 5 datasets were removed due to extreme values in GABA+ (*n* = 4) and Glx (*n* = 1) levels (Q3+3*IQR). Quality metrics and metabolite levels for the final sample are presented in Table 2, and the group average difference spectrum for each voxel and stimulus block is presented in Figure 3.

**Figure 3.**
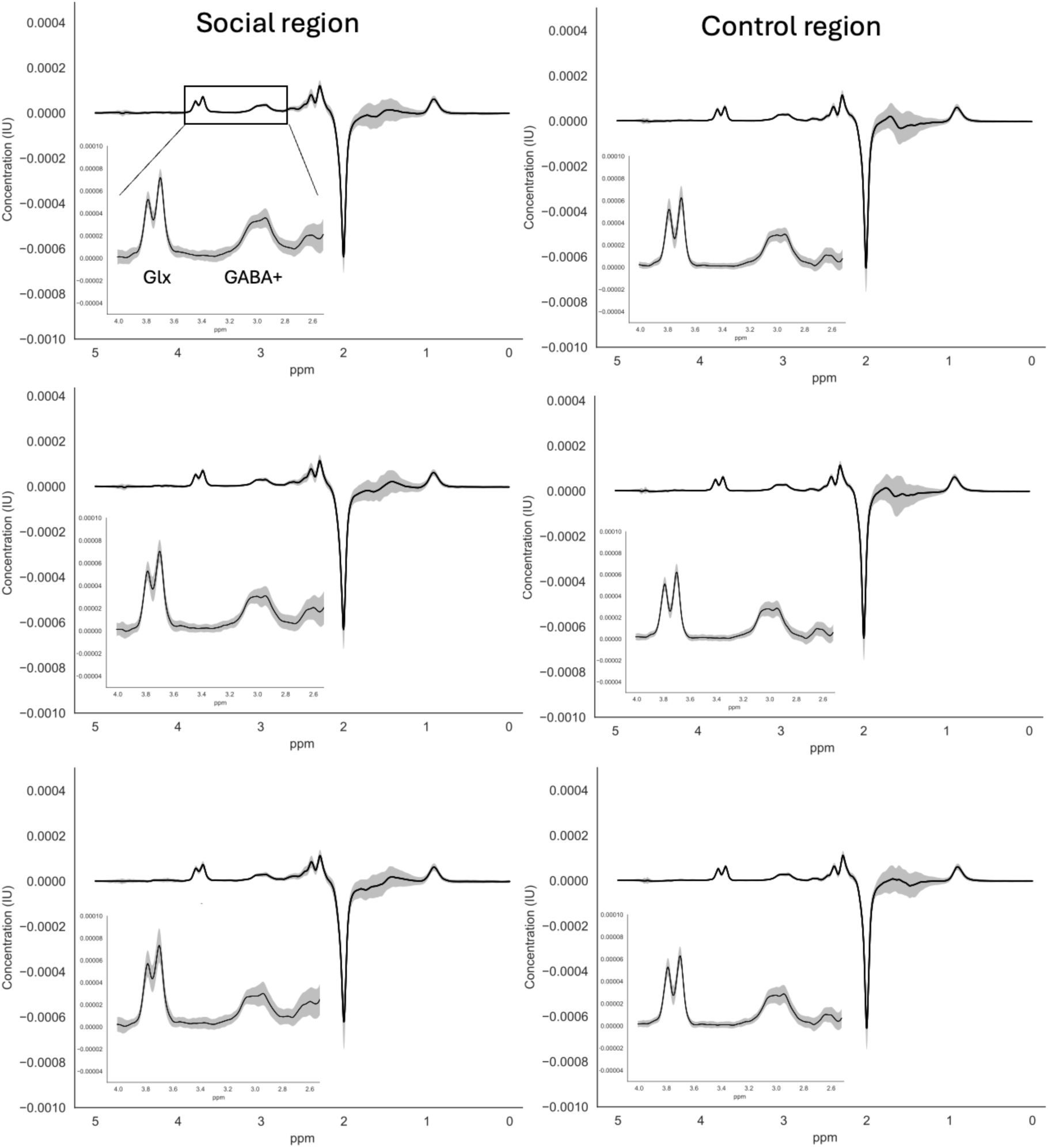
Average chemical shift spectrum. Average difference chemical shift spectrum across all participants for social brain (left) and control (right) voxels for the first rest block (top panel), first functional block (middle), and second functional block (bottom). Inset illustrates glutamate+glutamine (Glx) and GABA+macromolecules (GABA+) peaks. Shading represents the standard deviation of the difference spectrum. ppm = parts per million, i.u. = institutional units.

**Table 2.** Quality metrics and metabolite levels for each stimulus block and voxel.

|  | Social region |  |  | Control region |  |  |
| --- | --- | --- | --- | --- | --- | --- |
|  | M(SD) |  |  | M(SD) |  |  |
|  | Rest | Faces | CB | Rest | Faces | CB |
| Water FWHM (Hz) |  | 10.68(1.11) |  |  | 8.97(.26) |  |
| NAA FWHM (Hz) | 8.6(.98) | 8.8(1.06) | 8.9(1.12) | 7.7(.52) | 7.8(.54) | 7.7(.55) |
| GABA+ fit error (%) | 6.7(2.32) | 7.4(2.18) | 7.3(2.03) | 5.1(.99) | 5.2(1.18) | 5.2(1.14) |
| GABA+ SNR | 11.5(2.93) | 11.7(6.12) | 10.8(4.53) | 14.1(2.49) | 13.3(2.57) | 13.2(2.14) |
| Glx fit error (%) | 3.3(1.0) | 3.5(.95) | 3.4(.96) | 2.7(.44) | 2.6(.52) | 2.6(.46) |
| Glx SNR | 23.1(5.33) | 23.7(4.69) | 23.6(4.51) | 26.7(3.69) | 26.1(4.07) | 26.4(4.34) |
| GABA+ (i.u.) | 4.2(1.31) | 3.7(.88) | 3.3(.72) | 4.1(.51) | 4.0(.45) | 3.9(.40) |
| Glx (i.u.) | 13.9(1.77) | 14.6(2.64) | 14.7(2.21) | 12.9(1.33) | 13.0(1.61) | 13.5(1.38) |

Notes: M = mean, SD = standard deviation, CB = checkerboard, FWHM = full width-half maximum, SNR = signal to noise ratio, GABA+ = GABA + metabolites, Glx = glutamate+glutamine, i.u. = institutional units.

For the sliding time-window analysis, 60-TR windows sliding by 10 TRs yield 73 time-points per participant per voxel (total N = 2,920 data-points per voxel per metabolite). Data-points were excluded if spectral fitting failed to fit the data, resulting in nine excluded from social brain (GABA+ *n* = 2, Glx *n* = 7) and none excluded from the control region. The data were then assessed for quality using the following criteria: FWHM > 15Hz or GABA+ or Glx fit error > 15% for both voxels. The following data-points were removed based on quality criteria: social brain GABA+ *n* = 298, Glx *n* = 15; control region GABA *n* = 2, Glx *n* = 0. Finally, data- points with extreme metabolite concentrations exceeding Q3+3×IQR were excluded: social brain GABA+ *n* = 261, Glx *n* = 25; control region GABA *n* = 42, Glx *n* = 0. Final data-points used in the sliding time-window statistical analysis were: social brain GABA+ *n* = 2,360 (80.8% retained), Glx *n* = 2,874 (98.4% retained); control region GABA+ *n* = 2,876 (98.5% retained), Glx *n* = 2,920 (100% retained). Given the sliding window analysis tends to have lower SNR and thus higher fit error, we also ran the same analyses with a fit error threshold of 65% (Pasanta et al., 2025a)(see Supplementary Material).

### 3.2. fMRS results

#### 3.2.1. Block results

For GABA+, the mixed effects model showed no significant voxel by stimulus interaction (*F*(2, 181.85) = 2.39, *p* = .094; Figure 4a). There was a main effect for voxel (*F*(1, 184.39) = 19.42, *p* < .001), with GABA+ levels lower in the social brain than control region, and for stimulus (*F*(2, 182.69) = 5.16, *p* = .006), with post hoc tests showing GABA+ was reduced for checkerboard compared to resting state (*t*(184) = 3.52, *p* = .002).

**Figure 4.**
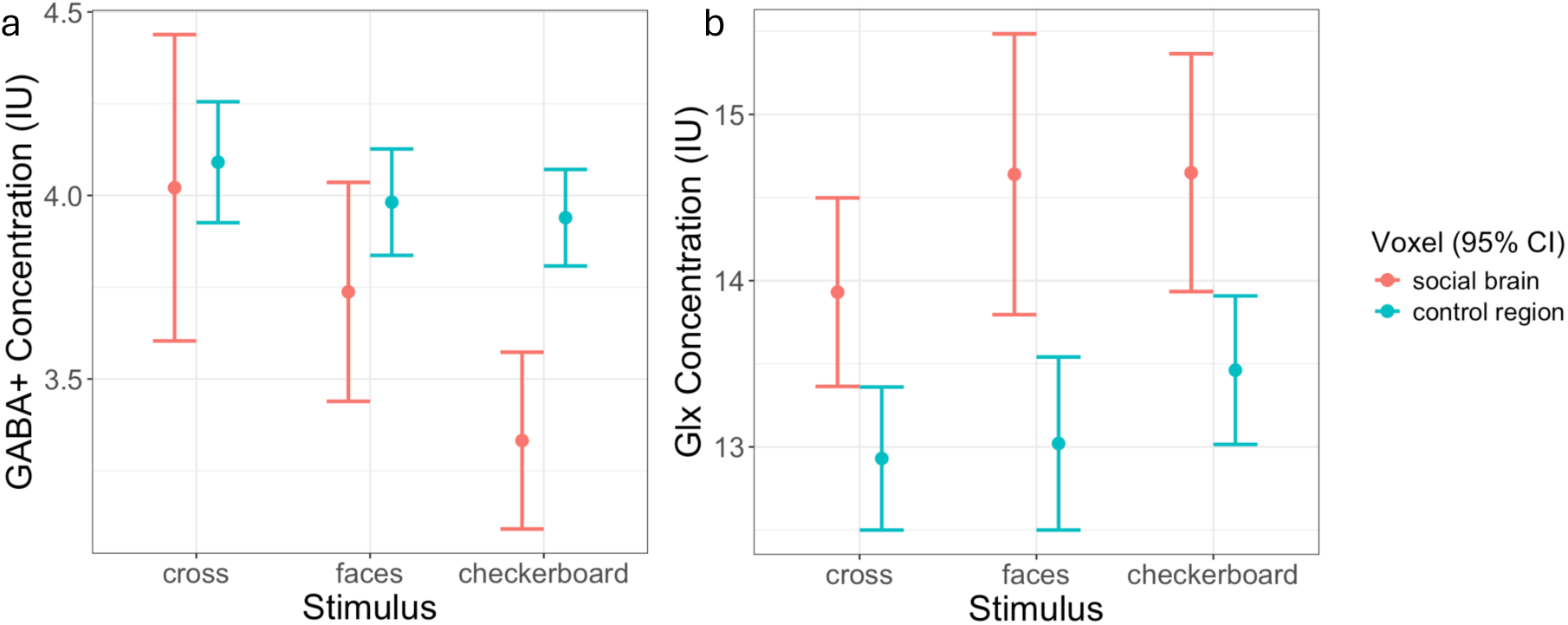
Plots of mean metabolite levels. Metabolite levels institutional units (IU) in social brain (red) and control (blue) voxels across stimulus types for a) GABA+ and b) glutamate+glutamine (Glx).

For Glx, there was also no significant voxel by stimulus interaction (*F*(2, 189.99) = .652, *p* = .522; Figure 4b). There was a significant main effect for voxel (*F*(1, 191.63) = 38.34, *p* < .001), with Glx levels higher in the social brain than control region, and for stimulus (*F*(2, 190.03) = 8.69, *p* < .001) with post hoc tests showing Glx was marginally higher (non-significant) for checkerboard than resting state (*t*(191) = -2.39, *p* = .053).

Follow-up Bayesian analyses were conducted within each voxel to examine the likelihood of the alternative hypothesis – difference between stimulus type. For social brain, there was moderate evidence for the alternative for GABA+ (BF: 3.29), with a .697 decrease in GABA+ for checkerboard versus resting state (95%CrI: [−1.137, −.257]), and a .283 decrease in GABA+ for faces versus resting state (95%CrI: [-0.738, 0.170]). There was no evidence for the alternative for Glx (BF: .24). For the control region, there was no evidence for the alternative for GABA+ (BF: .21) or Glx (BF: .28).

#### 3.2.2. Sliding time-window results

Dynamic changes in GABA+ and Glx levels over time for each participant are presented in Figure 5. Bayesian hierarchical models with an AR(1) autocorrelation term were fitted to examine condition effects on metabolite levels in social brain and control voxels with the first resting state block as baseline. Convergence diagnostics indicated satisfactory performance (all R-hat ≤ 1.01). Full model summaries and ROPE results are provided in Supplementary Table 1.

**Figure 5.**
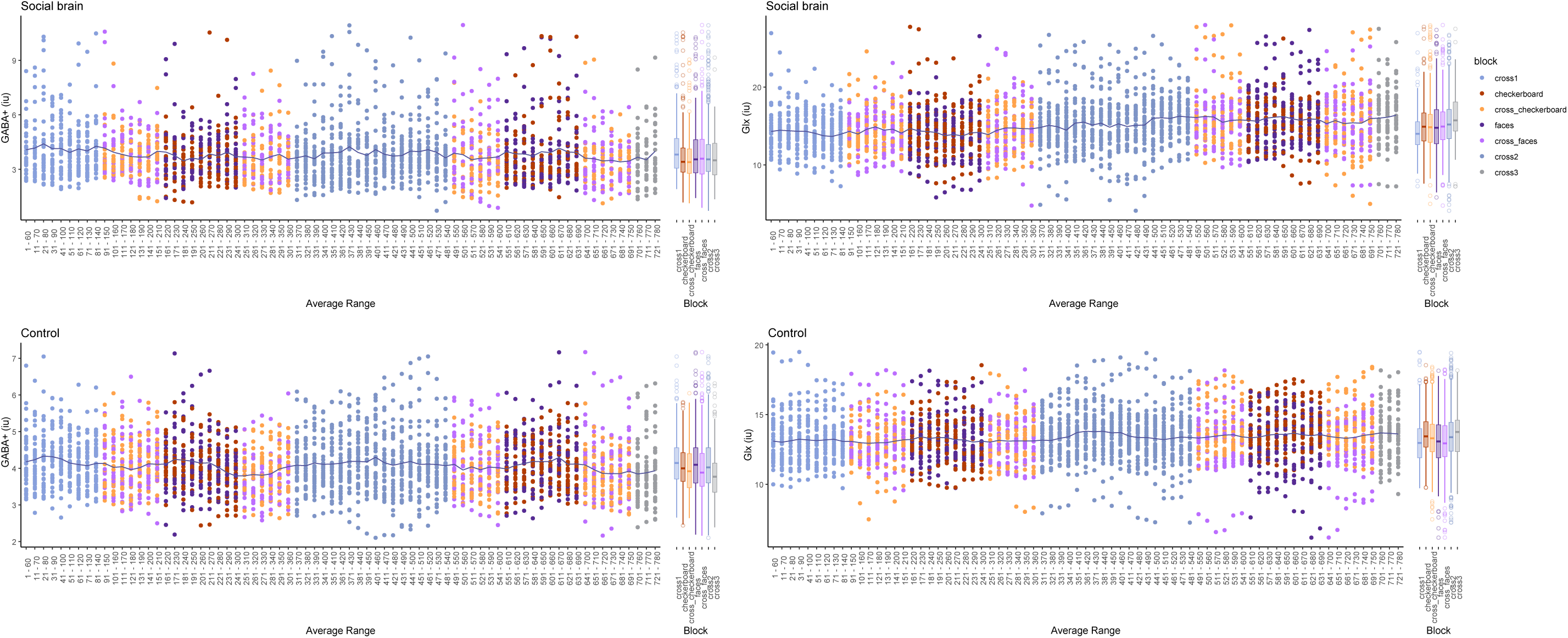
Sliding time-window dynamics. GABA+ (left) and Glx (right) concentration dynamics from sliding time-window analysis for social brain (top) and control regions (bottom). Average range represents the time at the centre of each analysis window (60 repetition times; 120 seconds). Colour coding indicates stimulus type: fixation cross (blue), dynamic faces (purple) and checkerboard (orange), with lighter colours indicating analysis of data from two stimulus types, e.g., light purple indicates analysis windows where fixation cross and dynamic faces are included. The line fit represents the mean value of metabolite levels within each analysis window. i.u. = institutional units.

In social brain, there was little evidence of GABA+ modulation for between conditions, with ROPE analysis suggesting no credible effects (Figure 6a). However, Glx in the social brain voxel demonstrated consistent reductions during time-windows included both functional and resting state blocks compared to the first resting state block. In particular, Glx decreased during social blocks (Median = -0.96, 95% CI [-1.76, -0.20], 0.0% inside ROPE), with ROPE analysis demonstrating this effect to be considered significant (<1% inside ROPE; Figure 6b).

**Figure 6.**
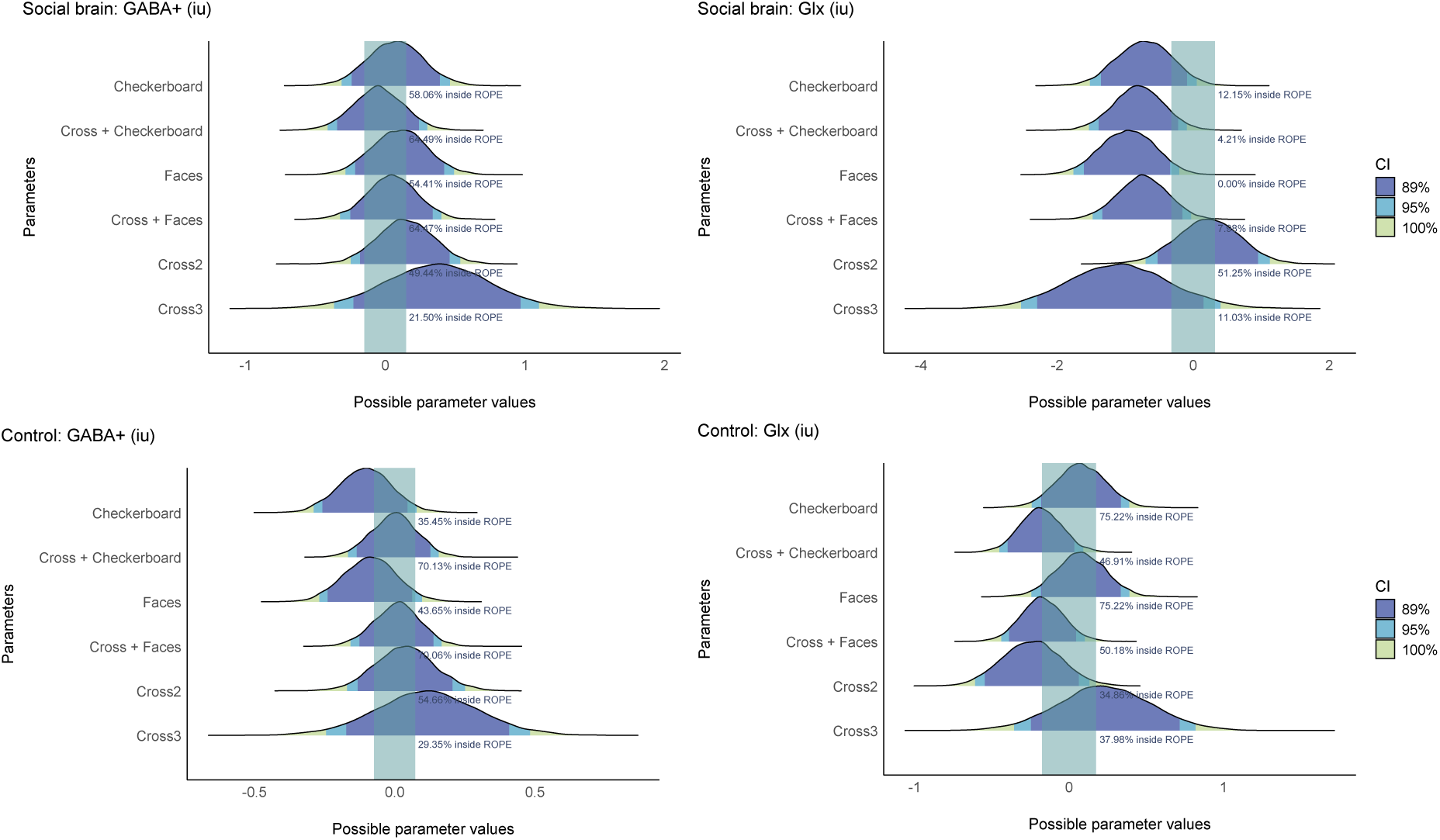
Range of Practical Equivalence (ROPE) analyses. ROPE analysis for the social brain GABA+ (a) and Glx (b), and control voxel GABA+ (c) and Glx (d). Shaded areas indicate posterior intervals (89%, 95%, 100%), with green boxes denoting the ROPE (±0.1 SD of the outcome, representing negligible effects). Values show the percentage of the 89% Highest Density Interval falling within the ROPE.

In the control region, there was little evidence for condition-related modulation of GABA+ or Glx. ROPE analysis similarly suggested no credible effects. Although weak trends suggested possible decreases in GABA+ under social and checkerboard stimulus presentation, and modest fluctuations in Glx, the 95% credible intervals all included zero, providing no firm evidence for systematic effects (Figure 6c and 6d).

#### 3.2.3. fMRS and social processes correlations

Higher SRS-2 scores were moderately correlated with higher levels of resting social brain Glx (rho = 0.325, *p* = .040, BF = 1.13, Figure 5), although did not survive FDR correction (*p_corrected_* = .243), no other correlations between SRS-2 scores and metabolite levels were significant (*p*s > .05). There were also no correlations between GABA+ and Glx across resting state and function tasks in either brain region (rho = -0.236 – 0.256, *p*s > 0.1). The full correlation table and scatterplots are presented in Supplementary Table 2 and Supplementary Figure 2.

### 3.3. Combined fMRS and fMRI results

fMRI analyses demonstrated significant BOLD signal in the expected brain regions for the faces and checkerboard stimuli (Figure 7). Compared to rest, the dynamic faces stimuli produced significant BOLD signal (*p*_corrected_ < .05) in left and right superior and middle temporal regions, TPJ, fusiform gyrus, amygdala, lateral occipital lobe, and prefrontal cortex, all of which are involved in social information processing, and validated our positioning of the social brain MRS voxel. The flashing checkerboard stimulus produced significant BOLD signal (*p*_corrected_ < .05), largely in the occipital cortex (Figure 7), validating our intended voxel placement, although only part of the significant BOLD signal produced by the checkerboard stimulus was captured within the control voxel, reflecting the constraints of voxel size and anatomical placement in visual cortex.

**Figure 7.**
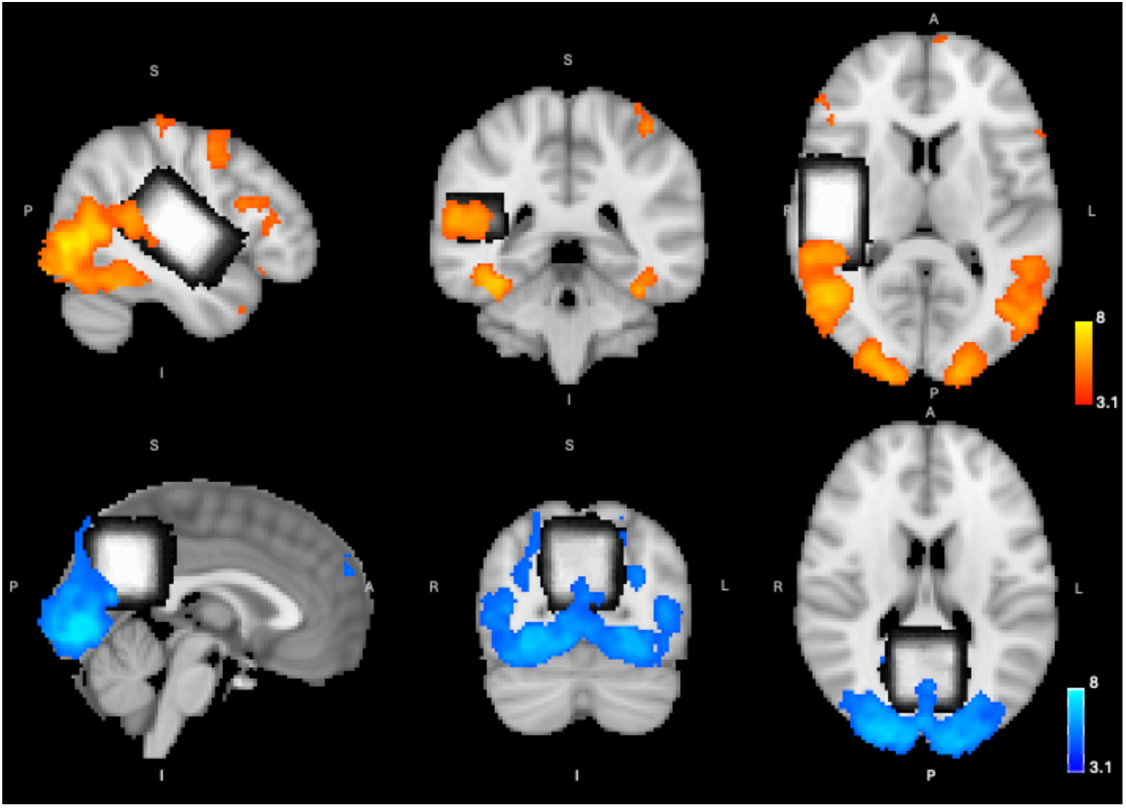
Group averaged mean fMRI bold signal. Group averaged fMRI BOLD signal to mean dynamic faces (top) and mean flashing checkerboard (bottom) compared to rest, with social brain (top) and control region (bottom) MRS voxels overlay. *Z*-statistic > 3.1, *p_corrected_* < .05.

The correlation between stimulus-related metabolite levels and BOLD signal was investigated by first creating a stimulus (faces or checkerboard) minus rest Glx and GABA value for each participants to align with mean BOLD signal (stimuli minus rest). There was a significant correlation between BOLD signal and Glx levels to the dynamic faces stimuli in the control region (*r* = 0.409, *p* = .010, *p*_corrected_ = .039, BF = 6.63), with BOLD signal increasing as Glx levels increased. No other correlations were significant, and no evidence for the alternative using Bayesian analyses. The full correlation table and scatterplots are presented in Supplementary Table 3 and Supplementary Figure 3.

## 4. Discussion

We investigated the modulating effect of dynamic social stimuli on Glx and GABA+ levels in a brain region associated with social information processing. We used a novel sliding time-window approach (Pasanta et al., 2024), as well as block analyses, to examine shifts in excitatory and inhibitory brain processes during social and non-social stimuli. Despite utilising a dynamic faces task known to elicit BOLD signal in right STG (Deen et al., 2015), the dynamic faces did not significantly modulate Glx or GABA+ levels in the right social brain voxel including STG. Similarly the flashing checkerboard stimulus did not modulate Glx or GABA+ in the control region (occipital cortex).

Analyses showed mixed results, with evidence for reduced inhibitory GABA+ in social brain based on block analyses, as well as reduced Glx in social brain based on sliding time-window analyses.

While conventional analyses showed no significant changes, Bayesian analyses provided evidence for reduced GABA+ during both dynamic faces and flashing checkerboard stimuli in the social brain while engaged in social and non-social stimulation. These findings may reflect changes in inhibitory processes within the STG/TJP, and/or insular cortex. However, as MRS-derived GABA+ reflects the total metabolite pool and does not distinguish between synaptic, intracellular, or extracellular compartments, changes in GABA+ cannot be interpreted as a direct index of inhibitory neurotransmission (Duncan et al., 2014; Mullins et al., 2014; Pasanta et al., 2023). The direction and underlying neurobiological mechanism of this change therefore remains uncertain. Reduced inhibitory processes is consistent with interpretations of previous research showing reduced GABA associated with stimulation, learning, and improved task performance (Heba et al., 2016; Kolasinski et al., 2019; Stagg et al., 2011). However, a recent meta-analysis demonstrated that GABA is not consistently modulated by sensory stimulation or task demands, potentially due to significant variability in data collection and analysis methods (Pasanta et al., 2023).

To improve sensitivity of our analyses and better understand stimulus-related metabolite dynamics, we implemented novel sliding time-window analyses in this study. This method allows for the quantification of small segments of spectroscopy data, which are then combined to demonstrate the progressive shift in metabolite levels. Sliding time-window analyses showed no meaningful differences in GABA+ levels for either functional task compared to rest (cross1) in either brain region, although there was a meaningful reduction in social brain Glx while participants viewed the dynamic faces stimuli. These findings differed from block analysis, likely reflecting differences in temporal resolution. Sliding window approaches may capture transient dynamics that are averaged in the block analysis, but are also more susceptible to increased noise and autocorrelation due to fewer transients per window (Ip & Bridge, 2022; Mullins, 2018; Pasanta et al., 2023, 2025b). Given the exploratory nature of the sliding window analysis, these results should be interpreted with appropriate caution.

Several methodological and neurobiological factors may explain the absence of robust task-related metabolite changes. First, it is possible that the intensity of the task was not enough to elicit a large enough shift in metabolites that would be detectable by MRS. Participants were asked to passively view a series of short videos that have previously elicited BOLD activity in STG (Deen et al., 2015). However, given the demand and social challenge of this task was low, the task itself may not have required the substantial shift in excitatory or inhibitory brain processes as predicted. Second, the very nature of edited MRS data collection requires a voxel of at least 27ml for adequate signal (Peek et al., 2023). This trade-off between voxel size (for SNR) and regional specificity in turn leads to the inclusion of additional non-social brain regions within the voxel. Placement of such a large voxel in the temporal region is also difficult due to proximity to sinuses, CSF, and skull. Furthermore, while MRS cannot determine the precise cellular or subcellular location of GABAergic and glutamatergic neurotransmitters (e.g., neuronal, synaptic, glial, intracellular, or extracellular), emerging evidence suggests that MR-visible signal may vary across compartments due to differences in molecular mobility and relaxation properties (Duncan et al., 2014). As such, task-related changes in MRS-detected metabolites may partly reflect shifts in compartmental distribution or MR visibility, rather than absolute changes in neurotransmitter concentration. This further complicates interpretation of fMRS findings in terms of underlying neurophysiological processes. Third, given we opted to acquire one continuous run of HERMES data with two resting state and two functional blocks in two separate brain regions, we faced a trade off between collecting enough data and participant tolerance. The default number of transients for a HERMES sequence is 320 (160 ONs, 160 OFFs) (Saleh et al., 2016), however, our paradigm included 150 transients for the first rest and two stimulus blocks, and 240 transients for the second rest block to account for metabolite lag following stimulus blocks. Given the data had good signal to noise during piloting, we opted for the shorter total scan time to reduce the likelihood of participant movement. It is possible though that the length of the scan blocks were too short to detect shifts in stimulus-related metabolite levels (Pasanta et al., 2023). Finally, the HERMES sequence is designed to specifically quantify GABA and GSH, with the combined signal of glutamate and glutamine (Glx) co-edited in the GABA-edited acquisition. Therefore, the Glx level is not a direct reflection of excitatory glutamate, given the contribution of glutamine, although edited Glx has been shown to correlate with specific quantifications of glutamate (Maddock et al., 2018).

These data show no significant correlation between Glx and GABA+ levels in either brain region, therefore we focus on changes in GABA+ and Glx separately rather than a Glx/GABA+ ratio. The lack of correlation between GABA+ and Glx may reflect differences in cellular composition within the voxel, and the use of a composite Glx measure, as discussed above (Rideaux et al., 2022). Other studies using editing sequences such as HERMES and MEGA-PRESS to quantify GABA+ as well as Glx have reported similarly low correlations (Rideaux, 2021; Rideaux et al., 2022; van Veenendaal et al., 2018). Future research could quantify glutamate more directly using a short-TE PRESS sequence or similar.

There were also no significant relationships between resting or stimulus-related GABA+ or Glx levels and RDoC Social Processes measured via the SRS-2, except for a moderate increase in resting social brain Glx levels with increased SRS-2 scores that did not survive FDR correction. The lack of associations between excitatory and inhibitory metabolites and autistic traits are in line with some clinical and neurotypical studies (Cochran et al., 2015; Demler et al., 2023), while others have shown significant correlations in neurotypical populations in similar brain regions (Ford et al., 2017a). This variability across studies may reflect differences in brain regions examined and quantification approaches. In relation to this study specifically, the suboptimal quantification of Glx via the HERMES editing sequence may explain the lack of relationships between Glx and social communication function.

In order to validate the dynamic faces task used herein, we acquired fMRI BOLD data on the identical faces and checkerboard stimuli. Significantly elevated BOLD signal was seen in the STG for faces, and occipital cortex for the checkerboard, as expected. We also examined whether fMRI BOLD signal in the MRS voxel locations was associated with GABA+ and Glx levels, finding only that, during the dynamic faces stimuli, increased BOLD signal was associated with increased Glx levels in the control brain region. Although the results must be considered preliminary, visual inspection of the corrected BOLD signal during the faces stimuli shows overlap with the control voxel. This could be explained by the location of the control voxel including precuneus, which is associated with social interactions (Petrini et al., 2014). Changes in BOLD activity during visual stimulation have been associated with increased glutamate and reduced GABA levels in occipital cortex (Bednařík et al., 2015; Ip et al., 2017, 2019; Koush et al., 2021; Muthukumaraswamy et al., 2009), while increased BOLD during a cognitive task has been associated with reduced GABA+ (Kühn et al., 2016) and increased Glx (Vijayakumari et al., 2018). However, others have found no associations between task-related glutamate or GABA+ and BOLD signal (Apšvalka et al., 2015; Costigan et al., 2019; Craven et al., 2023). In this study, no other associations between BOLD signal and stimulus-related metabolite levels were observed. This may be due to the relatively limited significant BOLD signal in the MRS voxels, or a lack of association between BOLD and Glx or GABA+ in these regions. These mixed findings highlight the need for further research to understand the relationship between BOLD signal and metabolites associated with excitation and inhibition during various functional tasks.

### 4.1. Limitations

The results of this study should be interpreted with consideration of some limitations. First, data were collected in the absence of eye or face monitoring, and although participants were responsive between blocks, it cannot be guaranteed they kept their eyes open for the entire scan. Second, a known limitation of edited MRS GABA quantification is the unavoidable inclusion of *J*-coupled macromolecular peaks that are co-edited with the GABA signal at 3ppm (Cudalbu et al., 2021; Mullins et al., 2014). Variability in individual macromolecular contributions can affect the specificity of the GABA signal detected and therefore blur the effect of stimulus on GABA levels. Furthermore, we used the HERMES spectral editing approach, which has a 4-step cycle (two for GABA), and thus limiting temporal resolution at the acquisition level, which may further contribute to the constrains described above. Third, as discussed above, the lack of effect of the flashing checkerboard stimulus on control region GABA+ may be due to suboptimal positioning voxel placement in visual cortex.

While the control voxel extended beyond the intended primary visual cortex region due to anatomical and shimming constraints, the resulting placement remained predominantly within visual cortex and was considered appropriate for indexing lower-level visual processing relative to the socially selective responses observed in STG/TPJ (Deen et al., 2015; Pitcher et al., 2011).

Nevertheless, partial inclusion of adjacent regions such as precuneus may have contributed to some overlap with social-task-related BOLD activity. Future studies could modify the shape of the voxel from cube to a rectangular prism for a more optimal fit. Nevertheless, several studies have found no effect of checkerboard in visual cortex (Bednařík et al., 2015; Dwyer et al., 2021; Kurcyus et al., 2018), suggesting variations in stimulus, acquisition techniques, or analysis methods contribute to study inconsistencies. Fourth, there was limited spatial overlap between the MRS voxel and peak task-related BOLD activation (see Figure 7). Group-level BOLD responses were observed to be more posterior and inferior relative to the mean voxel location, which may have introduced partial volume effects and reduced sensitivity to detect metabolite changes specific to the most strongly activated regions. Although voxel placement was optimised based on anatomical landmarks and prior literature, the inherent mismatch between functional activation peaks and the relatively large MRS voxel likely contributed to spatial averaging across functionally heterogeneous tissue.

Additionally, the block size of 60 TRs, while chosen to maximise temporal resolution, does not constitute a complete phase cycle for the HERMES acquisition scheme. Although no systematic subtraction artefacts were observed in the present data, future fMRS studies employing HERMES should adopt block sizes that are integer multiples of the full phase cycling scheme (e.g., 64 TRs) to ensure complete coherence cancellation. Here we also did not explicitly account for potential line- narrowing BOLD effect (Bednařík et al., 2015; Just, 2024), future fMRS studies could benefit from incorporating linewidth-linewidth matching task-related T2* changes to spectrum analysis. Fifth, although spectral registration and quality control analyses indicated that scanner drift in this data were corrected for, subtle temporal effects across the relatively long acquisition may still have contributed to block-order-related variability. These effects could include small residual frequency instabilities, participant habituation or fatigue, or other slow physiological changes occurring over time (Mullins, 2018; Pasanta et al., 2023; Rideaux, 2020). Such factors may be particularly relevant in fMRS paradigms where relatively subtle metabolite changes are being examined across extended acquisitions. Finally, as noted above, MRS does not allow localisation of metabolites to specific cellular or synaptic compartments limiting interpretation of underlying neurobiological mechanisms (Duncan et al., 2014).

### 4.2. Conclusions

This study probed shifts in GABA+ and Glx levels in social brain regions in response to socially laden stimuli. Although block analyses shoed no significant changes in GABA+ and Glx, Bayesian analyses showed some evidence for decreased GABA+ in social brain during both the checkerboard and social stimulus. Sliding time-window analysis, however, showed a reduction in social brain Glx during social stimuli. Future studies could incorporate a more challenging social task with measurable engagement to investigate changes in social brain Glx and GABA+ and examine alternate social brain regions such as medial prefrontal cortex. Nevertheless, while modest, results demonstrate feasibility into the application of fMRS in social and cognitive neuroscience.

## Acknowledgements

The authors acknowledge the facilities and scientific and technical assistance of the National Imaging Facility, a National Collaborative Research Infrastructure Strategy (NCRIS) capability, at the Swinburne Neuroimaging (SNI) Facility, Swinburne University of Technology. This study was funded by a Swinburne Neuroimaging Access and Support Scheme Enhanced Capability Grant (2002- TFDW). NP is supported by a Simons SFARI Human Cognitive and Behavioral Science award. NP is supported through the MRC Centre for Neurodevelopmental Disorders. This study represents independent research in part funded by the National Institute for Health and Care Research (NIHR) Maudsley Biomedical Research Centre (BRC) at South London and Maudsley NHS Foundation Trust and King’s College London.

## Data availability

Study related code can be found at https://osf.io/y7jqp/. Raw MRI data is available on OpenNeuro (https://openneuro.org/datasets/ds007806).

## Notes

https://openneuro.org/datasets/ds007806

https://osf.io/y7jqp/

### Competing Interest Statement

The authors have declared no competing interest.

### Summary of Updates

Reviews conducted buy request from peer-reviewers, including additional clarifications on a number of data collection and analysis decision, and re-analysis of the sliding time-window data at a more stringent threshold (exclusion fit error < 15%).

