## Supplementary Material for "Examining excitation-inhibition modulation during social processing using functional Magnetic Resonance Spectroscopy"

**Supplementary Figure 1:** Functional MRI task diagram. Note: faces stimuli de-identified.

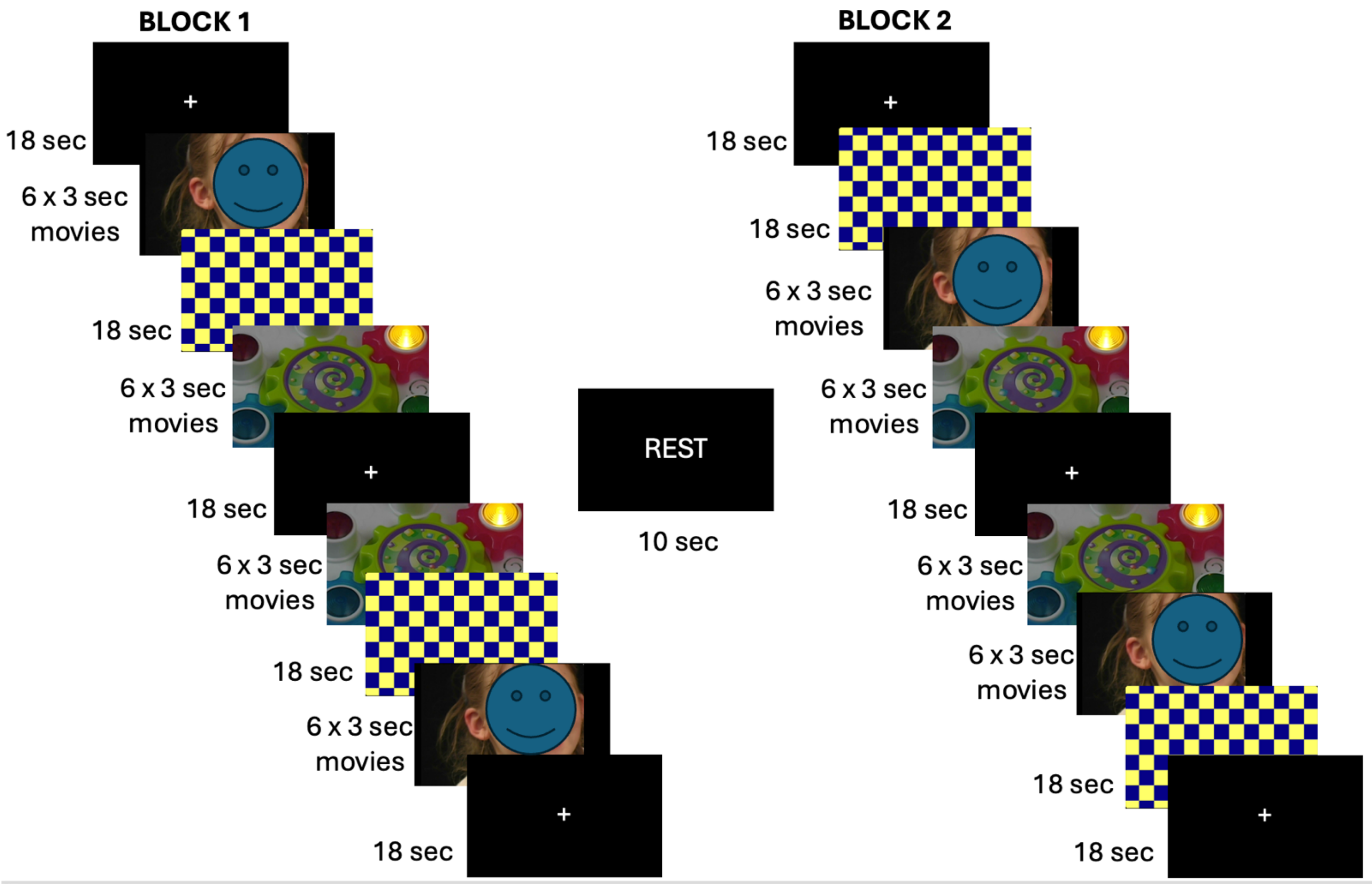


### fmriprep analysis description

Results included in this manuscript come from preprocessing performed using fMRIPrep 25.1.3 (Esteban et al. (2019); Esteban et al. (2018); RRID:SCR_016216), which is based on Nipype 1.10.0 (K. Gorgolewski et al. (2011); K. J. Gorgolewski et al. (2018); RRID:SCR_002502).

#### Preprocessing of B0 inhomogeneity mappings

A total of 1 fieldmaps were found available within the input BIDS structure for each subject. A B0 nonuniformity map (or fieldmap) was estimated from the phase-drift map(s) measure with two consecutive GRE (gradient-recalled echo) acquisitions. The corresponding phase-map(s) were phase-unwrapped with prelude (FSL None).

#### Anatomical data preprocessing

A total of two T1-weighted (T1w) images were found within the input of each participants BIDS dataset. Each T1w image was corrected for intensity non-uniformity (INU) with N4BiasFieldCorrection (Tustison et al. 2010), distributed with ANTs 2.6.2 (Avants et al. 2008, RRID:SCR_004757). The T1w-reference was then skull-stripped with a Nipype implementation of the antsBrainExtraction.sh workflow (from ANTs), using OASIS30ANTs as target template. Brain tissue segmentation of cerebrospinal fluid (CSF), white-matter (WM) and gray-matter (GM) was performed on the brain-extracted T1w using fast (FSL (version unknown), RRID:SCR_002823, Zhang, Brady, and Smith 2001). An anatomical T1w-reference map was computed after registration of 2 <module ‘nipype.interfaces.image’ from ‘/opt/conda/envs/fmriprep/lib/python3.12/site-packages/nipype/interfaces/image.py’> images (after INU-correction) using mri_robust_template (FreeSurfer 7.3.2, Reuter, Rosas, and Fischl 2010). Brain surfaces were reconstructed using recon-all (FreeSurfer 7.3.2, RRID:SCR_001847, Dale, Fischl, and Sereno 1999), and the brain mask estimated previously was refined with a custom variation of the method to reconcile ANTs-derived and FreeSurfer-derived segmentations of the cortical gray-matter of Mindboggle (RRID:SCR_002438, Klein et al. 2017).

Volume-based spatial normalization to the MNI152NLin6Asym standard spaces was performed through nonlinear registration with antsRegistration (ANTs 2.6.2), using brain-extracted versions of both T1w reference and the T1w template (FSL’s MNI ICBM 152 non-linear 6th Generation Asymmetric Average Brain Stereotaxic Registration Model [Evans et al. (2012), RRID:SCR_002823; TemplateFlow ID: MNI152NLin6Asym).

**Functional data preprocessing**

For each participant’s BOLD run, the following preprocessing was performed. First, a reference volume was generated, using a custom methodology of fMRIPrep, for use in head motion correction. Head-motion parameters with respect to the BOLD reference (transformation matrices, and six corresponding rotation and translation parameters) were estimated before any spatiotemporal filtering using mcflirt (FSL , Jenkinson et al. 2002). The estimated fieldmap was then aligned with rigid-registration to the target EPI (echo-planar imaging) reference run. The field coefficients were mapped on to the reference EPI using the transform. The BOLD reference was then co-registered to the T1w reference using bbregister (FreeSurfer) which implements boundary-based registration (Greve and Fischl 2009). Co-registration was configured with six degrees of freedom. Several confounding time-series were calculated based on the preprocessed BOLD: framewise displacement (FD), DVARS and three region-wise global signals. FD was computed using two formulations following Power (absolute sum of relative motions, Power et al. (2014)) and Jenkinson (relative root mean square displacement between affines, Jenkinson et al. (2002)). FD and DVARS are calculated for each functional run, both using their implementations in Nipype (following the definitions by Power et al. 2014). The three global signals are extracted within the CSF, the WM, and the whole-brain masks. Additionally, a set of physiological regressors were extracted to allow for component-based noise correction (CompCor, Behzadi et al. 2007). Principal components are estimated after high-pass filtering the preprocessed BOLD time-series (using a discrete cosine filter with 128s cut-off) for the two CompCor variants: temporal (tCompCor) and anatomical (aCompCor). tCompCor components are then calculated from the top 2% variable voxels within the brain mask. For aCompCor, three probabilistic masks (CSF, WM and combined CSF+WM) are generated in anatomical space. The implementation differs from that of Behzadi et al. in that instead of eroding the masks by 2 pixels on BOLD space, a mask of pixels that likely contain a volume fraction of GM is subtracted from the aCompCor masks. This mask is obtained by dilating a GM mask extracted from the FreeSurfer’s aseg segmentation, and it ensures components are not extracted from voxels containing a minimal fraction of GM. Finally, these masks are resampled into BOLD space and binarized by thresholding at 0.99 (as in the original implementation). Components are also calculated separately within the WM and CSF masks. For each CompCor decomposition, the k components with the largest singular values are retained, such that the retained components’ time series are sufficient to explain 50 percent of variance across the nuisance mask (CSF, WM, combined, or temporal). The remaining components are dropped from consideration. The head-motion estimates calculated in the correction step were also placed within the corresponding confounds file. The confound time series derived from head motion estimates and global signals were expanded with the inclusion of temporal derivatives and quadratic terms for each (Satterthwaite et al. 2013). Frames that exceeded a threshold of 0.9 mm FD or 1.5 standardized DVARS were annotated as motion outliers. Additional nuisance timeseries are calculated by means of principal components analysis of the signal found within a thin band (crown) of voxels around the edge of the brain, as proposed by (Patriat, Reynolds, and Birn 2017). All resamplings can be performed with a single interpolation step by composing all the pertinent transformations (i.e. head-motion transform matrices, susceptibility distortion correction when available, and co-registrations to anatomical and output spaces). Gridded (volumetric) resamplings were performed using nitransforms, configured with cubic B-spline interpolation.

### Statistical analysis

Statistical models: the full analysis pipeline is available at osf.io/

#### Mixed Linear Models

```
model_GABA <- lmer(log(GABA) ~ voxel*stimulus + block_order + (1|PID), data=data, REML=TRUE, na.action = na.ex
clude)

model_Glx <- lmer(Glx ~ voxel*stimulus + block_order + (1|PID), data=data, REML=TRUE, na.action = na.exclude)
```

#### Bayesian Models

```
model_GABA_STS <- brm(GABA ~ stimulus + (1|PID), data = sts_data, family = gaussian(), prior = c(
  prior(normal(0, 5), class = "b"),
  prior(normal(0, 5), class = "Intercept"),
  prior(student_t(3, 0, 5), class = "sd")
 ), chains = 4, cores = 4, iter = 2000, silent = TRUE)


sts_GABA <- sts_data[!is.na(sts_data$GABA),]

models <- anovaBF(formula = GABA ~ stimulus, data = sts_GABA)
```

**Supplementary Table 1.** Bayesian model results from the social brain and control regions for each stimulus type compared to baseline cross1. For the QA threshold of fit error of 15%.

| Parameter | Median | 95% CI | pd | ROPE | %in ROPE | R-hat | ESS |
|---|---|---|---|---|---|---|---|
| **Social brain: GABA+** | | | | | | | |
| Intercept | 3.8 | [ 3.49, 4.11] | 100% | [-0.15, 0.15] | 0% | 1.002 | 3166 |
| Checkerboard | 0.07 | [-0.31, 0.46] | 63.86% | [-0.15, 0.15] | 58.06% | 1.001 | 3543 |
| Fixation cross (rest) + Checkerboard | -0.05 | [-0.41, 0.30] | 61.61% | [-0.15, 0.15] | 64.49% | 1.001 | 3320 |
| Social stimuli | 0.1 | [-0.29, 0.49] | 69.73% | [-0.15, 0.15] | 54.41% | 1.001 | 3523 |
| Fixation cross (rest) + Social stimuli | 0.05 | [-0.33, 0.40] | 60.81% | [-0.15, 0.15] | 64.47% | 1.001 | 3282 |
| Fixation cross2 (rest2) | 0.14 | [-0.25, 0.54] | 76.25% | [-0.15, 0.15] | 49.44% | 1 | 4313 |
| Fixation cross3 (rest3) | 0.37 | [-0.37, 1.10] | 83.97% | [-0.15, 0.15] | 21.50% | 1.001 | 5549 |
| **Social brain: Glx** | | | | | | | |
| Intercept | 15.47 | [14.74, 16.22] | 100% | [-0.32, 0.32] | 0.00% | 1 | 3834 |
| Checkerboard | -0.71 | [-1.51, 0.05] | 96.59% | [-0.32, 0.32] | 12.15% | 1 | 4795 |
| Fixation cross (rests) + Checkerboard | -0.8 | [-1.53, -0.09] | 98.58% | [-0.32, 0.32] | 4.21% | 1 | 4490 |
| Social stimuli | -0.96 | [-1.76, -0.20] | 99.44% | [-0.32, 0.32] | 0.00% | 1 | 4746 |

| | | | | | | | |
|---|---|---|---|---|---|---|---|
| Fixation cross (rests) + Social stimuli | -0.74 | [-1.48, -0.03] | 97.94% | [-0.32, 0.32] | 7.98% | 1 | 4550 |
| Fixation cross (rest2) | 0.22 | [-0.70, 1.12] | 68.76% | [-0.32, 0.32] | 51.25% | 1 | 6000 |
| Fixation cross (rest3) | -1.09 | [-2.53, 0.40] | 92.09% | [-0.32, 0.32] | 11.03% | 1 | 7496 |
| **Control region: GABA+** | | | | | | | |
| Intercept | 4.1 | [ 3.94, 4.26] | 100% | [-0.07, 0.07] | 0% | 1 | 3686 |
| Checkerboard | -0.1 | [-0.29, 0.08] | 86.96% | [-0.07, 0.07] | 35.45% | 1.001 | 3969 |
| Fixation cross (rests) + Checkerboard | -3.49E-04 | [-0.17, 0.16] | 50.10% | [-0.07, 0.07] | 70.13% | 1.001 | 3826 |
| Social stimuli | -0.09 | [-0.27, 0.10] | 82.70% | [-0.07, 0.07] | 43.65% | 1.001 | 3967 |
| Fixation cross (rests) + Social stimuli | 0.01 | [-0.16, 0.17] | 55.29% | [-0.07, 0.07] | 70.06% | 1.001 | 3746 |
| Fixation cross2 (rest2) | 0.04 | [-0.17, 0.25] | 63.51% | [-0.07, 0.07] | 54.66% | 1.001 | 5761 |
| Fixation cross3 (rest3) | 0.12 | [-0.25, 0.48] | 74.25% | [-0.07, 0.07] | 29.35% | 1 | 6872 |
| **Control region: Glx** | | | | | | | |
| Intercept | 13.36 | [12.85, 13.85] | 100% | [-0.18, 0.18] | 0% | 1.003 | 1054 |
| Checkerboard | 0.08 | [-0.24, 0.39] | 68.55% | [-0.18, 0.18] | 75.22% | 1 | 2817 |
| Fixation cross (rests) + Checkerboard | -0.18 | [-0.45, 0.09] | 90.90% | [-0.18, 0.18] | 46.91% | 1.001 | 2858 |
| Social stimuli | 0.08 | [-0.24, 0.39] | 68.81% | [-0.18, 0.18] | 75.22% | 1 | 2821 |
| Fixation cross (rests) + Social stimuli | -0.17 | [-0.44, 0.10] | 89.72% | [-0.18, 0.18] | 50.18% | 1 | 2876 |
| Fixation cross2 (rest2) | -0.24 | [-0.61, 0.13] | 89.92% | [-0.18, 0.18] | 34.86% | 1 | 4721 |
| Fixation cross3 (rest3) | 0.23 | [-0.36, 0.82] | 78.15% | [-0.18, 0.18] | 37.98% | 1 | 6867 |

Notes: CI = confidence interval; pd = probability direction; ROPE = Range of Practical Equivalence; ESS = effective sample size; GABA+ = GABA + macromolecules; Glx = glutamate + glutamine.

**Supplementary Table 2:** Spearman rank correlations between Social Responsiveness Scale scores and GABA+ and Glx concentrations by voxel and stimulus type.

| | GABA+ | | | Glx | | |
|---|---|---|---|---|---|---|
| | rho | *p*-value | *p*-FDR | rho | *p*-value | *p*-FDR |
| **Social brain** | | | | | | |
| Cross | 0.01 | 0.974 | 0.974 | 0.33 | 0.040* | 0.243 |
| Social stimuli | 0.05 | 0.791 | 0.974 | 0.09 | 0.584 | 0.583 |
| Checkerboard | 0.02 | 0.918 | 0.974 | 0.16 | 0.329 | 0.395 |
| **Control region** | | | | | | |
| Cross | -0.08 | 0.619 | 0.974 | 0.21 | 0.209 | 0.313 |
| Social stimuli | 0.22 | 0.177 | 0.531 | 0.25 | 0.128 | 0.257 |
| Checkerboard | 0.30 | 0.067 | 0.400 | 0.28 | 0.089 | 0.256 |

Notes: FDR = corrected for false discovery rate; GABA+ = GABA + macromolecules; Glx = glutamate + glutamine.

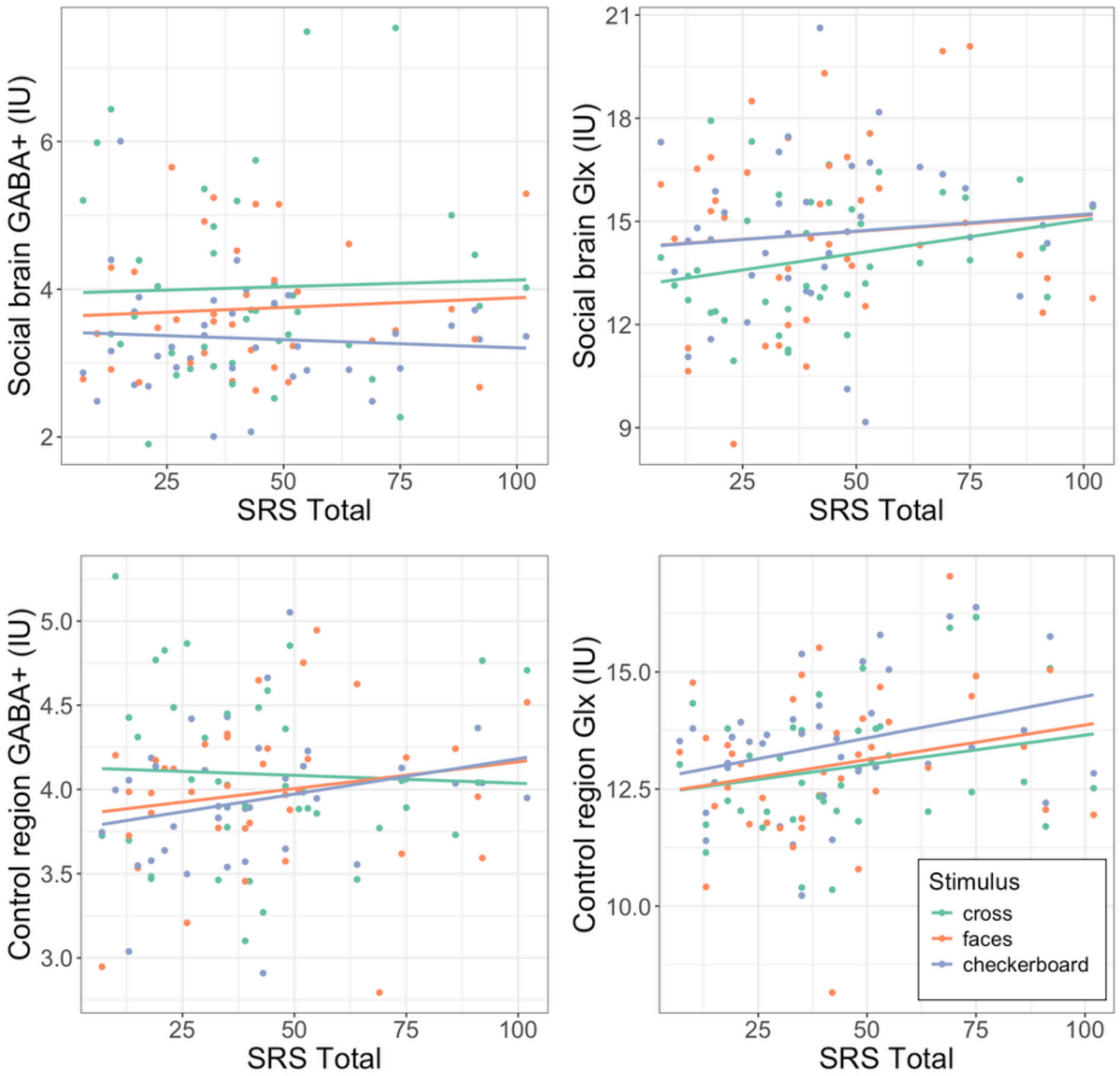


**Supplementary Figure 2.** Scatterplots of social brain and control region GABA+ and Glx concentrations associations with Social Responsiveness Scale (SRS) scores. IU = institutional units.

**Supplementary Table 3.** Pearsons *r* correlations between social brain and control voxel GABA+ and Glx concentrations and mean BOLD signal in equivalent brain regions for dynamic faces and flashing checkerboard stimuli minus baseline (fixation cross).

| | GABA+ | | | Glx | | |
|---|---|---|---|---|---|---|
| | *r* | *p*-value | *p*-FDR | *r* | *p*-value | *p*-FDR |
| **Social brain** | | | | | | |
| Faces | -0.038 | 0.830 | 0.830 | -0.080 | 0.626 | 0.626 |
| Checkerboard | 0.122 | 0.470 | 0.830 | -0.240 | 0.142 | 0.236 |
| **Control region** | | | | | | |
| Faces | -0.072 | 0.661 | 0.830 | 0.409 | 0.010* | 0.039* |
| Checkerboard | -0.084 | 0.618 | 0.830 | 0.221 | 0.177 | 0.236 |

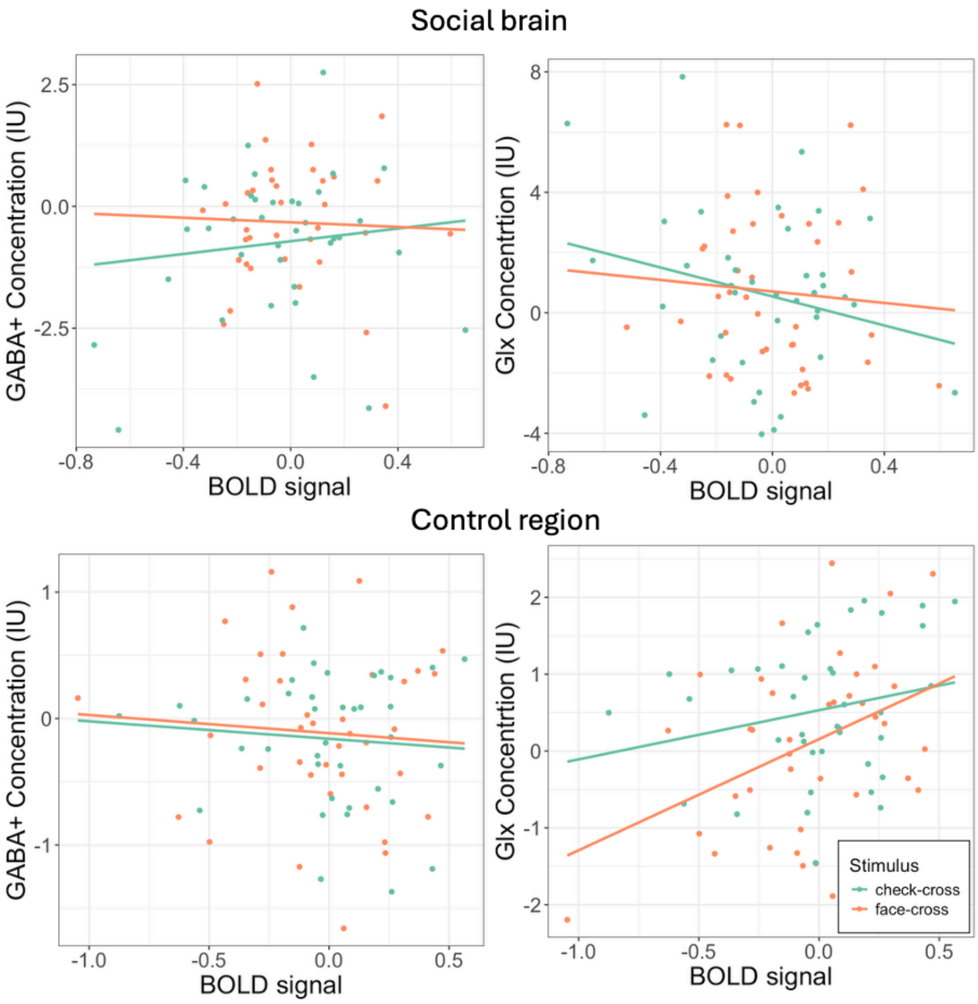


**Supplementary Figure 3.** Scatterplots of metabolite vs BOLD associations. Associations between mean BOLD signal (stimulus minus rest) and equivalent GABA+ and Glx concentration (stimulus minus cross) in the social brain region (top) and control region (bottom). IU = institutional units.

**Sliding time-window analysis with 65% fit error threshold**

Given the sliding window analysis tends to have lower SNR and thus higher fit error, we also ran the same analyses with a fit error threshold of 65% (Pasanta et al., 2025a). No data-points from either voxel were excluded based on these quality criteria. Data-points with extreme metabolite concentrations exceeding Q3+3×IQR were excluded. For the social brain voxel, 296 data-points were removed (GABA+ *n* = 263; Glx *n* = 28). For the control voxel, 42 data-points were removed (GABA+ *n* = 42, Glx *n* = 0). Final data-points were: social brain GABA+ *n* = 2,655 (91.0% retained), Glx *n* = 2,885 (98.8% retained); control region GABA+ *n* = 2,878 (98.6% retained), Glx *n* = 2,920 (100% retained).

**Supplementary Table 4.** Bayesian model results from the social brain and control regions for each stimulus type compared to baseline cross1. For the QA threshold of fit error of 65%.

| Parameter | Median | 95% CI | pd | ROPE | %in ROPE | R-hat | ESS |
|---|---|---|---|---|---|---|---|
| **Social brain: GABA+** | | | | | | | |
| Intercept | 3.55 | [ 3.24, 3.87] | 100% | [-0.15, 0.15] | 0% | 1.001 | 4482 |
| Checkerboard | 0.04 | [-0.34, 0.42] | 57.31% | [-0.15, 0.15] | 62.09% | 1 | 4495 |
| Fixation cross (rests) + Checkerboard | 0.18 | [-0.18, 0.54] | 83.46% | [-0.15, 0.15] | 43.15% | 1.001 | 4098 |
| Social stimuli | 0.00 | [-0.38, 0.39] | 50.16% | [-0.15, 0.15] | 62.25% | 1 | 4544 |
| Fixation cross (rests) + Social stimuli | 0.32 | [-0.05, 0.67] | 95.58% | [-0.15, 0.15] | 15.49% | 1.001 | 4138 |
| Fixation cross2 (rest2) | 0.05 | [-0.35, 0.46] | 59.67% | [-0.15, 0.15] | 59.52% | 1 | 5704 |
| Fixation cross3 (rest3) | 0.72 | [ 0.04, 1.42] | 98.05% | [-0.15, 0.15] | 0% | 1 | 8870 |
| **Social brain: Glx** | | | | | | | |
| Intercept | 15.49 | [14.74, 16.22] | 100% | [-0.32, 0.32] | 0% | 1 | 3830 |
| Checkerboard | -0.80 | [-1.60, -0.01] | 97.56% | [-0.32, 0.32] | 8.13% | 1 | 4525 |
| Fixation cross (rests) + Checkerboard | -0.98 | [-1.68, -0.25] | 99.71% | [-0.32, 0.32] | 0% | 1 | 4580 |
| Social stimuli | -0.65 | [-1.44, 0.15] | 94.21% | [-0.32, 0.32] | 17.79% | 1 | 4552 |
| Fixation cross (rests) + Social stimuli | -0.89 | [-1.60, -0.17] | 99.33% | [-0.32, 0.32] | 0.62% | 1 | 4564 |
| Fixation cross (rest2) | 0.21 | [-0.72, 1.12] | 67.36% | [-0.32, 0.32] | 52.82% | 1 | 5611 |
| Fixation cross (rest3) | -1.08 | [-2.61, 0.45] | 91.46% | [-0.32, 0.32] | 12.78% | 1 | 8037 |
| **Control region: GABA+** | | | | | | | |
| Intercept | 13.33 | [12.83, 13.83] | 100% | [-0.17, 0.17] | 0% | 1.002 | 941 |
| Checkerboard | 0.11 | [-0.19, 0.41] | 75.99% | [-0.17, 0.17] | 68.40% | 1 | 2930 |
| Fixation cross (rests) + Checkerboard | -0.17 | [-0.44, 0.11] | 88.74% | [-0.17, 0.17] | 51.15% | 1 | 3098 |
| Social stimuli | 0.09 | [-0.22, 0.39] | 71.09% | [-0.17, 0.17] | 74.62% | 1 | 2997 |
| Fixation cross (rests) + Social stimuli | -0.15 | [-0.42, 0.13] | 85.01% | [-0.17, 0.17] | 58.89% | 1 | 3054 |
| Fixation cross2 (rest2) | -0.18 | [-0.59, 0.21] | 80.95% | [-0.17, 0.17] | 49.68% | 1 | 4692 |
| Fixation cross3 (rest3) | 0.27 | [-0.33, 0.87] | 81.34% | [-0.17, 0.17] | 33.38% | 1 | 6672 |
| **Control region: Glx** | | | | | | | |
| Intercept | 4.11 | [ 3.94, 4.27] | 100% | [-0.07, 0.07] | 0% | 1.002 | 3527 |

| | | | | | | | |
|---|---|---|---|---|---|---|---|
| Checkerboard | -0.13 | [-0.31, 0.05] | 91.40% | [-0.07, 0.07] | 26.54% | 1.002 | 3423 |
| Fixation cross (rests) + Checkerboard | 0.02 | [-0.15, 0.19] | 58.53% | [-0.07, 0.07] | 65.77% | 1.001 | 3739 |
| Social stimuli | -0.14 | [-0.33, 0.04] | 93.85% | [-0.07, 0.07] | 20.69% | 1.002 | 3477 |
| Fixation cross (rests) + Social stimuli | 0.02 | [-0.15, 0.18] | 58.73% | [-0.07, 0.07] | 65.66% | 1.001 | 3754 |
| Fixation cross2 (rest2) | 0.01 | [-0.20, 0.23] | 55.16% | [-0.07, 0.07] | 55.70% | 1.001 | 4646 |
| Fixation cross3 (rest3) | 0.13 | [-0.22, 0.49] | 75.79% | [-0.07, 0.07] | 27.23% | 1 | 7380 |

Notes: CI = confidence interval; pd = probability direction; ROPE = Range of Practical Equivalence; ESS = effective sample size; GABA+ = GABA + macromolecules; Glx = glutamate + glutamine.
